# Structural and functional impact of non-synonymous SNPs in the CST complex subunit TEN1: Structural genomics approach

**DOI:** 10.1101/507046

**Authors:** Mohd. Amir, Vijay Kumar, Taj Mohammad, Ravins Dohare, Md. Tabish Rehman, Mohamed F. Alajmi, Afzal Hussain, Faizan Ahmad, Md. Imtaiyaz Hassan

## Abstract

TEN1 protein is a key component of CST complex, implicated in maintaining the telomere homeostasis, and provide stability to the eukaryotic genome. Mutations in *TEN1* gene have higher chances of deleterious impact; thus, interpreting the number of mutations and their consequential impact on the structure, stability and function is essentially important. Here, we have investigated the structural and functional consequences of nsSNPs in the *TEN1* gene. A wide array of sequence- and structure-based computational prediction tools were employed to identify the effects of 78 nsSNPs on the structure and function of TEN1 protein and deleterious nsSNPs were identified. These deleterious or destabilizing nsSNPs are scattered throughout the structure of TEN1. However, major mutations were observed in the α1-helix (12-16) and β5-strand (88-96). We further observed that mutations at C-terminal region were have higher tendency to form aggregate. In-depth structural analysis of these mutations reveals that the pathogenecity of these mutations are driven mainly through larger structural changes because of alterations in non-covalent interactions. This work provides a blue print to pinpoint the possible consequences of pathogenic mutations in the CST complex subunit TEN1.

## Introduction

Telomeres are consist of non-coding ends of eukaryotic linear chromosomes, and play a vital role in the replication, regulation and protection of genome [1,2]. Ends of eukaryotic chromosomes can be identified by recombination and repair system of the cells as DNA strand breaks, that often proceed to end-to-end fusion and instability of genome [3,4]. Telomeres along with telomere accessory complexes, such as shelterin and CST, suppress unwanted DNA damage response (DDR) and act as buffer between crucial genomic information and maintenance of chromosomes end [5,6]. Shelterin complex is composed of six subunits (TRF1, TRF2, RAP1, TIN2, TPP1 and POT1) which are located primarily to single- and double-stranded telomeric DNA [7]. In addition to repressing DDR and chromosome fusion, shelterin complex also caps the telomeric ends by facilitating the formation of T-loop. It is also acting as processivity factor via recruiting telomerase to chromosomes ends [8,9].

CST complex is consists of three subunits (CTC1, STN1 and TEN1), specifically localizes to the single stranded DNA (ssDNA) of telomere and is involved in telomere capping and regulation of telomere length (Chen et al., 2012; Nakaoka et al., 2012; Wellinger, 2009). However, increasing evidences demonstrated that the STN1-TEN1 complex has some extra telomeric functions as it is involved in resolving replication fork stalling during replication stress [10,11]. CST complex is also involved in the removal of G-quadruplexes (G4: G-rich repeats) [12]. The G-rich region of telomere is very prone to form G4 throughout telomeric DNA and poses severe challenges for telomere replication machinery [13]. In addition, CST complex binds to the 3’ ends of telomeres and regulates polymerase α-mediated syntheses of C-strand [14].

In addition to polymerase α-mediated syntheses of C-strand, a subunit of CST (CTC1-STN1) regulates telomerase mediated extension of G-rich overhang which is critical for the cell proliferation. Deficiency of CTC1-STN1 complex leads to overextension of G-rich overhangs which initiate DDR [15,16]. In this process, the role of TEN1 is indispensable as it is essential to provide stability to CTC1-STN1 complex. Disruption of TEN1 results in progressive shortening of telomere more like caused by telomerase deficiency.As telomere maintenance is paramount to genome stability, mutations in the genes encoding essential components of CST are associated with varieties of genetic abnormalities including cancer [17], Coat plus [18,19,20] and Dyskeratosis Congenita [21,22].

Prediction of nsSNPs affecting protein structure and function in detail may be investigated by the aid of cutting-edge computational methods. In many cases, nsSNPs have little or no effect on protein structure and functions, but sometime a single mutation is highly lethal [23]. Experimental studies suggested that about one-third of nsSNPs are deleterious to human health [24]. Thus, identification of such deleterious nsSNPs is of serious concern in terms of diagnosis and therapeutics prospective. A little in-silico large scale mutational analysis has been performed on nsSNPs of TEN1 protein [25]. Taking this opportunity into consideration and fact that TEN1 plays crucial role in the telomere maintenance, we have predicted the structural and functional effects of about 78 nsSNPs in the coding region of *TEN1* gene. The present study will offer indepth understanding of the role of nsSNPs on the structure and function of TEN1 protein.

## Materials and methods

### Data collection

Distribution of nsSNPs in human *TEN1* gene was retrieved from dbSNP [26], Ensembl [27] and HGMD [28] databases. Data enrichment was carried out by removing the variant duplicates of different databases. The human TEN1 amino acid sequence was obtained in FASTA format from UniProt database (UniProt ID: Q86WV5) (http://www.uniprot.org/). Three-dimensional (3D) structure of TEN1 (PDB ID: 4JOI) was downloaded from Protein Data Bank (PDB) [29]. Functional annotations of all SNPs were extracted from dbSNP database, for example whether the SNPs present in an intron or exon, in the 3’ or 5’ untranslated region (UTR), or downstream or upstream of the *TEN1* gene.

### Sequence based prediction of deleterious nsSNPs

SIFT (Sorting Intolerant from Tolerant) (http://sift.jcvi.org/) algorithm was used to predict the amino acid substitution as tolerable and intolerable depending upon the physical and sequence-homology features. Substitutions with normalized probabilities of ≥0.05 and ≤0.05 were predicted as tolerated and deleterious, respectively [30,31]. There were about 78 nsSNPs identified from Ensembl and dbSNP databases. Prediction of tolerated and deleterious effect of these nsSNPs in human *TEN1* gene was predicted using SIFT. PROVEAN (Protein Variation Effect Analyzer) (http://provean.jcvi.org/) tool was used to predict the consequences of amino acid substitution on protein function [32]. It predicts nsSNPs as “deleterious” if the score is less than threshold value (cutoff is −2.5), and “neutral” if the predicted score is more than the cutoff value. All the nsSNPs in human *TEN1* gene was calculated and analyzed using this cutoff value.

PolyPhen-2 (Polymorphism Phenotyping-2) (http://genetics.bwh.harvard.edu/pph2/) was used to calculate functional predictions of coding variants. It uses a particular empirical rule comprises of both comparative and physical considerations to predict the probable functional impacts of mutation on the structure-function relationship. FASTA format of protein sequence was used as input to calculate the effects of a particular substitution [33]. It calculates position-specific independent count (PSIC) score for each substitution and then estimate the score deviations. A mutation is considered as possibly destructive mutation if the PSIC score is ≥0.9.

### Structure-based prediction of destabilizing nsSNPs

STRUM (https://zhanglab.ccmb.med.umich.edu/STRUM/) tool was used to predict the stability differences between Wt and mutant proteins. Initially, from protein sequences, a 3D model was generated by I-TASSER simulation and used to train STRUM model through gradient boosting regression. STRUM predicts the possible effects of nsSNPs on the structure and function of protein using conservation score from alignment of multiple-threading template. The query sequence used as input in FASTA format and calculates the impact of a particular substitution in a given sequence [34]. SDM2 (Site Direct Mutator 2) (http://structure.bioc.cam.ac.uk/sdm2) is a knowledge-based tool used to estimate the impact of mutations on stability of protein [35]. It uses constrained environment-specific substitution tables (ESSTs) to calculate the differences in the protein stability upon mutation [35,36]. SDM2 uses PDB as input file, and a point variants to estimates the stability difference score between the Wt and mutants.

PoPMuSiC (http://babylone.ulb.ac.be/popmusic/) tool was used to predict changes in thermodynamic stability upon mutation. PoPMuSiC employing a linear combination of statistical potentials whose coefficients depends on the solvent accessibility of the substituted residues. It uses PDB as input file. DUET server was used to predict the impact of mutations on the stability of TEN1 protein using PDB code. DUET calculate a combined or consensus predictions of SDM and mCSM (mutation Cutoff Scanning Matrix) using Support Vector Machines (SVMs) in a non-linear regression fashion. The output it provides is in the form of change in Gibbs free energy (ΔΔ*G*), where negative sign indicate destabilizing mutation [37]. mCSM was implicated to predict the impact of mutations on stability of proteins using graph-based structural signatures. It predicts protein-protein and protein-nucleic acid interaction [38].

### Identification of diseased phenotype

MutPred (http://mutpred.mutdb.org/) tool was used to predict the association of nsSNPs with disease phenotype [39]. It employs several attributes associated to structure, function and evolution using PSI-BLAST [40], SIFT [30] and Pfam profiles [41] together with structure disorder prediction tools such as TMHMM [42], DisProt [43] and MARCOIL [44]. Score with g-value more than 0.75 and p-value less than 0.05 are considered as confident hypothesis. PhD-SNP (http://snps.biofold.org/phd-snp/phd-snp.html) is online SVM based prediction tool, was used to predict the pathological effects of a given mutation [45].

### Aggregation propensity analysis

SODA (Protein **so**lubility from **d**isorder and **a**ggregation propensity) was used to predict the change in protein solubility upon mutation by comparing sequence profile of WT and mutants. The aggregation or intrinsic disorder score obtained from PASTA [46], and ESpritz [47], and a combined result obtained from Kyte-Doolittle [48] and FELLS [49]. SODA also predicts types of variation, including insertion and deletion in a given sequence [50].

### Sequence conservation analysis

The importance of a particular amino acid in the structure and functions of protein can be generally from its conservation score using multiple sequence alignment. The blue-print of amino acid conservation was identified by ConSurf tool which measures the degree of conservation of each amino acid at particular position along with evolutionary profile of amino acid sequence [51]. Conservation score was ranges from 1 to 9, where 1 depicts rapidly evolving (variable), 5 indicate region which are evolving moderately, and 9 shows slowly evolving (evolutionarily conserved) position. Exposed residues with high conservation score are being considered as functional whereas buried residues with high conservation score are believed as structural residues.

### Analysis of solvent accessibility

Relative side-chain solvent accessibility (RSA), residue depth and residue-occluded packing density (OSP) of Wt and mutant TEN1 protein have been performed using SDM2 server [35]. It uses ESSTs table to calculate the differences in their RSA, residue depth and OSP of Wt and mutant proteins. RSA have been calculated based on Lee and Richards method [52]. Three classes of relative RSA were defined based on the method of Lee and Richards, whereby a probe of given radius is rolled around the surface of the molecule [52].

## Results and Discussion

All reported SNPs of *TEN1* gene was extracted from Ensembl (http://www.ensembl.org/) and dbSNP databases (http://www.ncbi.nlm.nih.gov/snp). A total of about 5712 SNPs were mapped and classified into 9 different functional classes. Four major classes of SNPs in *TEN1* gene are shown in the **Fig. 1.** About 5250 SNPs were mapped in intronic region and approximately 78 was found in the coding non-synonymous/missense region. 5’ – and 3’ UTR-regions have 277 and 91 SNPs, respectively. In addition, 61 SNPs in coding synonymous, 5 SNPs in frame shift, 3 SNPs in each 3’ and 5’ splice site region are also observed. The present study focuses only on missense mutations mapped in the coding region. A total of 78 nsSNPs were taken for further analysis.

**Fig. 1:**
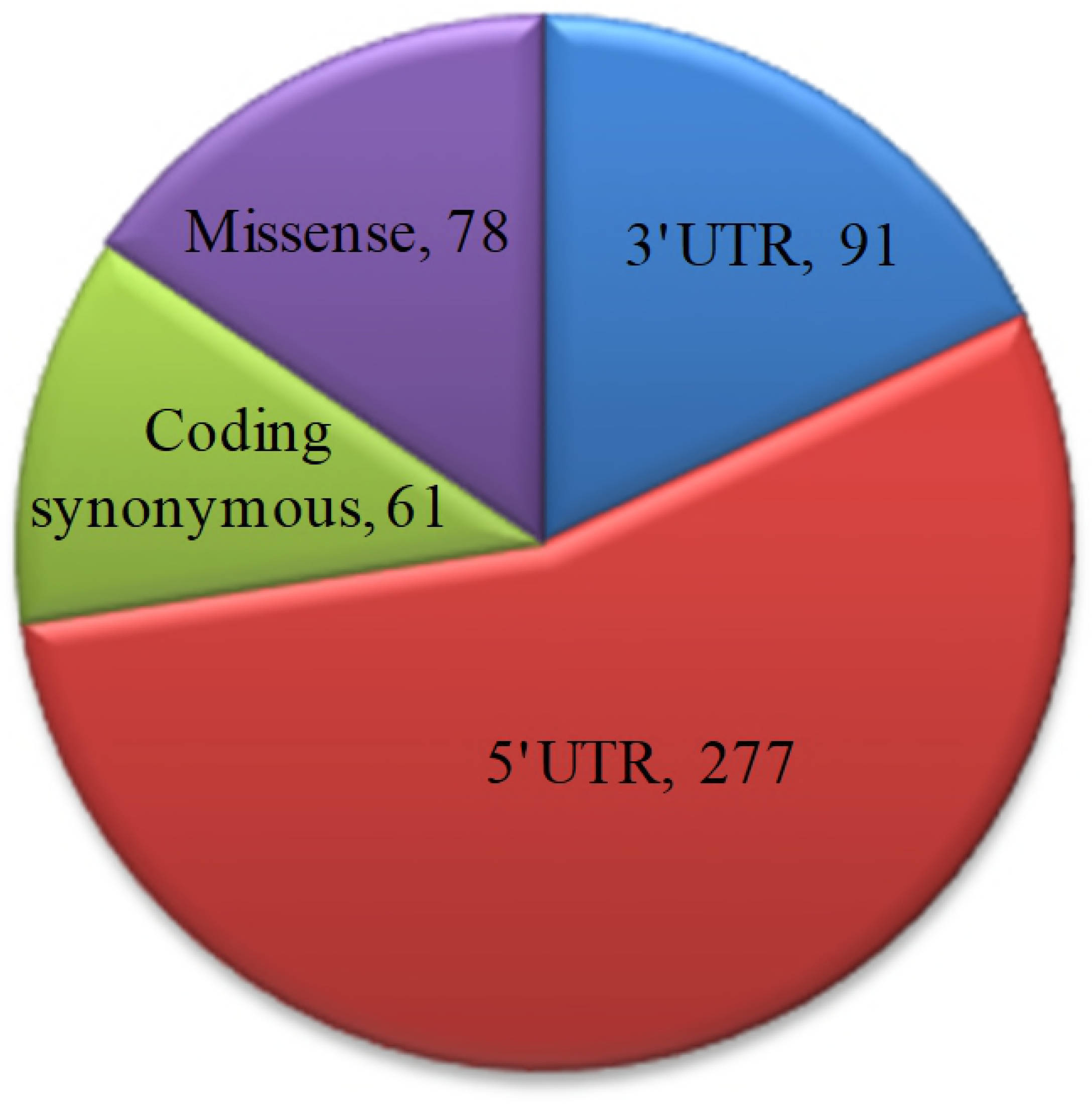
Pie chart representation of SNPs in *TEN1* gene using dbSNP database.

To identify the structural and functional impact on missense mutations in *TEN1* gene, we have employed a multi-tier approach. To collect high confidence nsSNPs in the *TEN1* gene, all mapped TEN1 nsSNPs were first subjected to sequence-based prediction using PolyPhen-2, PROVEAN and SIFT, followed by structure-based stability predictions using PoPMuSiC, SDM2, DUET, mCSM and STRUM web-servers. Further, distributions of high confidence nsSNPs were analyzed on the basis of their structure descriptors and phenotypic association. In consistence, we discuss pathogenic mutations in relation to their sequence conservation, functional importance and aggregation propensities. Finally, we expand our analysis and extensively analyzed the structural and functional impact of pathogenic mutations on the local environment of the TEN1 protein. An overview of computational methods used in this study is depicted in **Fig. 2**.

**Fig. 2:**
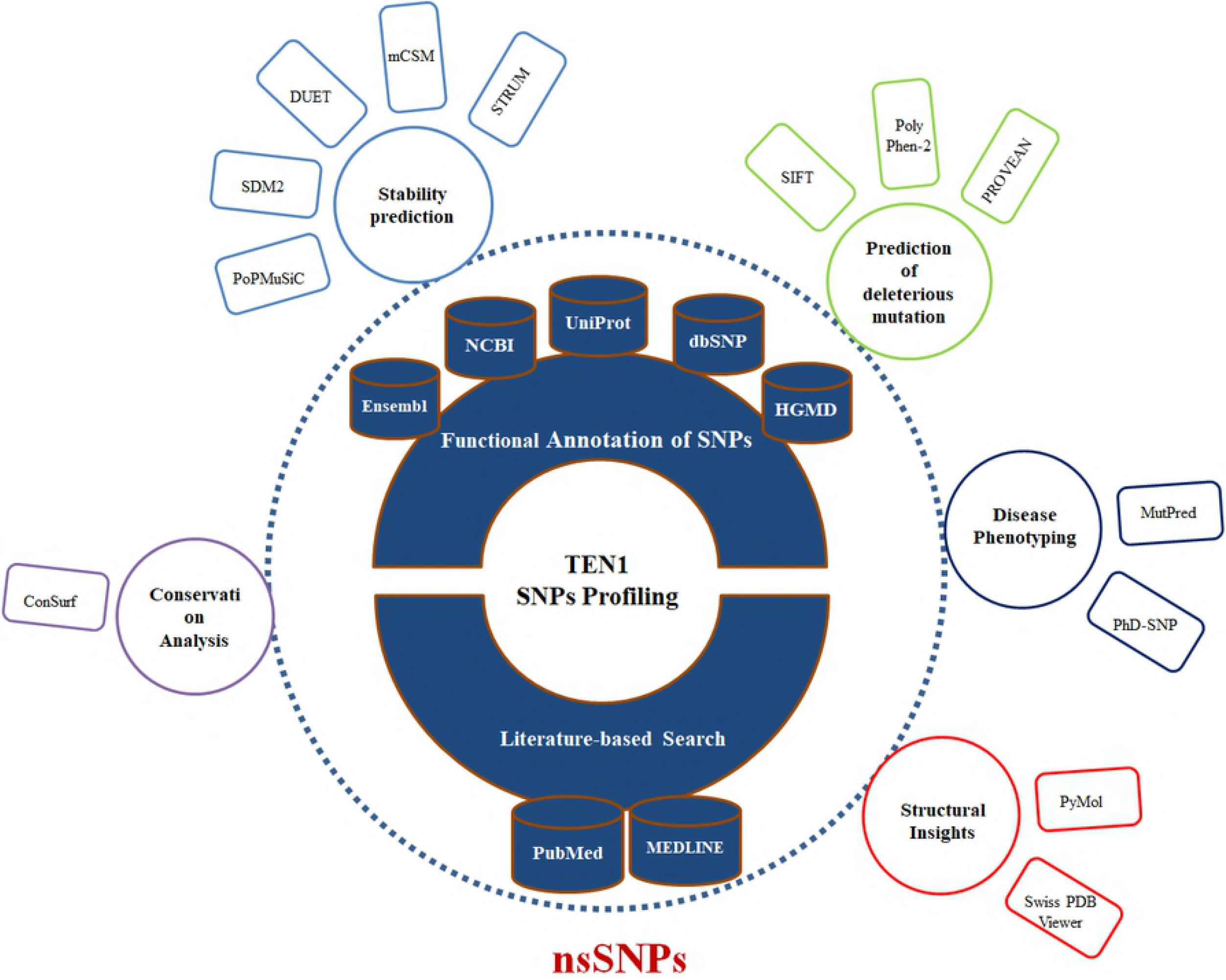
Overview of computational approaches used to identify the deleterious or pathogenic mutations in the TEN1 protein at structural and functional level.

### Identification of deleterious nsSNPs

To pinpoint the structural and functional consequences of nsSNPs in *TEN1* gene, we have performed an extensive structural analysis. The reason for using multiple tools is to improve the confidence level of prediction. Accumulation of deleterious nsSNPs using single approach may not always be satisfactory as some mutations that have score very close to cut-off value are prone to false prediction. Therefore, using multiple tools in both sequence- and structure-based predictions may provide an accurate result. nsSNPs predicted to be deleterious in at least two methods from sequence-based prediction methods and three tools depict destabilizing effects from structure-based prediction were collected and termed as “high confidence nsSNPs”.

Sequence-based prediction of all nsSNPs in *TEN1* gene was calculated by SIFT, PROVEAN and PolyPhen-2. A total of 78 nsSNPs of human *TEN1* gene was considered for analysis. Sequence- and structure-based predictions are listed in the **Table S1 and S2**. SIFT, PolyPhen-2 and PROVEAN predicted that out of 78 nsSNPs, 40 (51%), 42 (53%), 36 (46%) nsSNPs, respectively were deleterious (**Fig. 3**). Similarly, STRUM, mCSM, DUET, SDM2, and PoPMuSiC predicted that 40 (51%), 70 (89%), 62 (79%) 60 (76%) and 58 (74%) nsSNPs, respectively, as protein destabilizing (**Fig. 3**). We have further focused only on those mutations which are predicted to be deleterious and identified 34 mutations showing a destabilizing behavior.

**Fig. 3:**
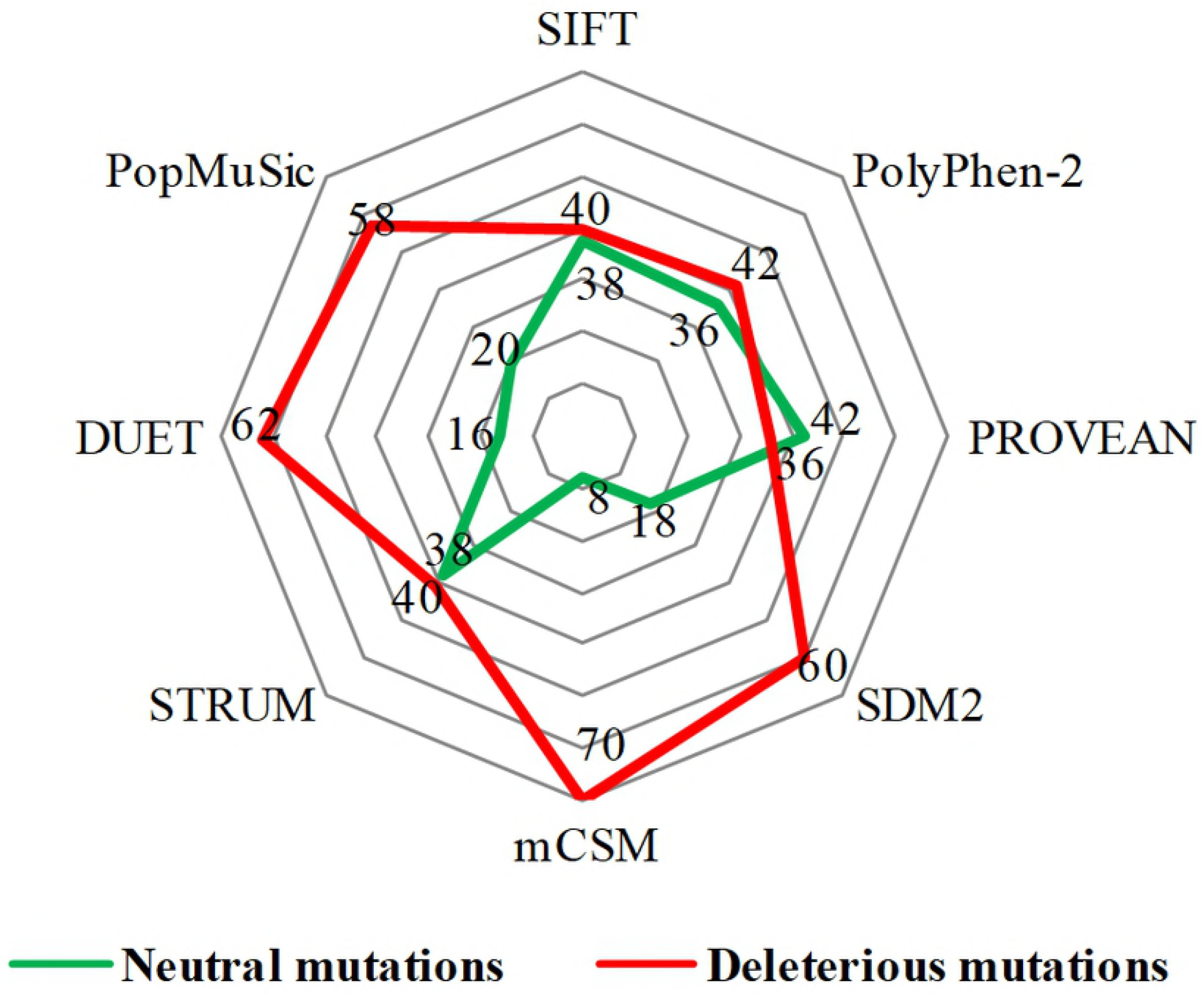
Distribution of predicted deleterious (red) and neutral (green) nsSNPs in *TEN1* gene.

### Sequence conservation analysis

A relative analysis of amino acid residue conservation based on protein sequence provides an understanding of the significance of particular amino acid residue and reveals its localized evolution. ConSurf results indicate that the amino acid residues stretch ranges, 26-32, 62-65, 75-78 and 91-99 were highly conserved (**Fig. 4**). The stretches of amino acids residues range, 32-61and 100-121 are not conserved. Further, structure-based conservation analysis suggested that amino acid residue belong to β1 (25-36) and L1-2 (37-40) (loop connecting β1 and β2), β4 (72-80) and β5 (88-96) are more conserved than β2 (41-48) and β3 (51-58) of TEN1 protein. Among these structural components, β5 (88-96) is the highly conserved while L4-5 (81-87) (loop connecting β4 and β5) is the least conserved.

**Fig. 4:**
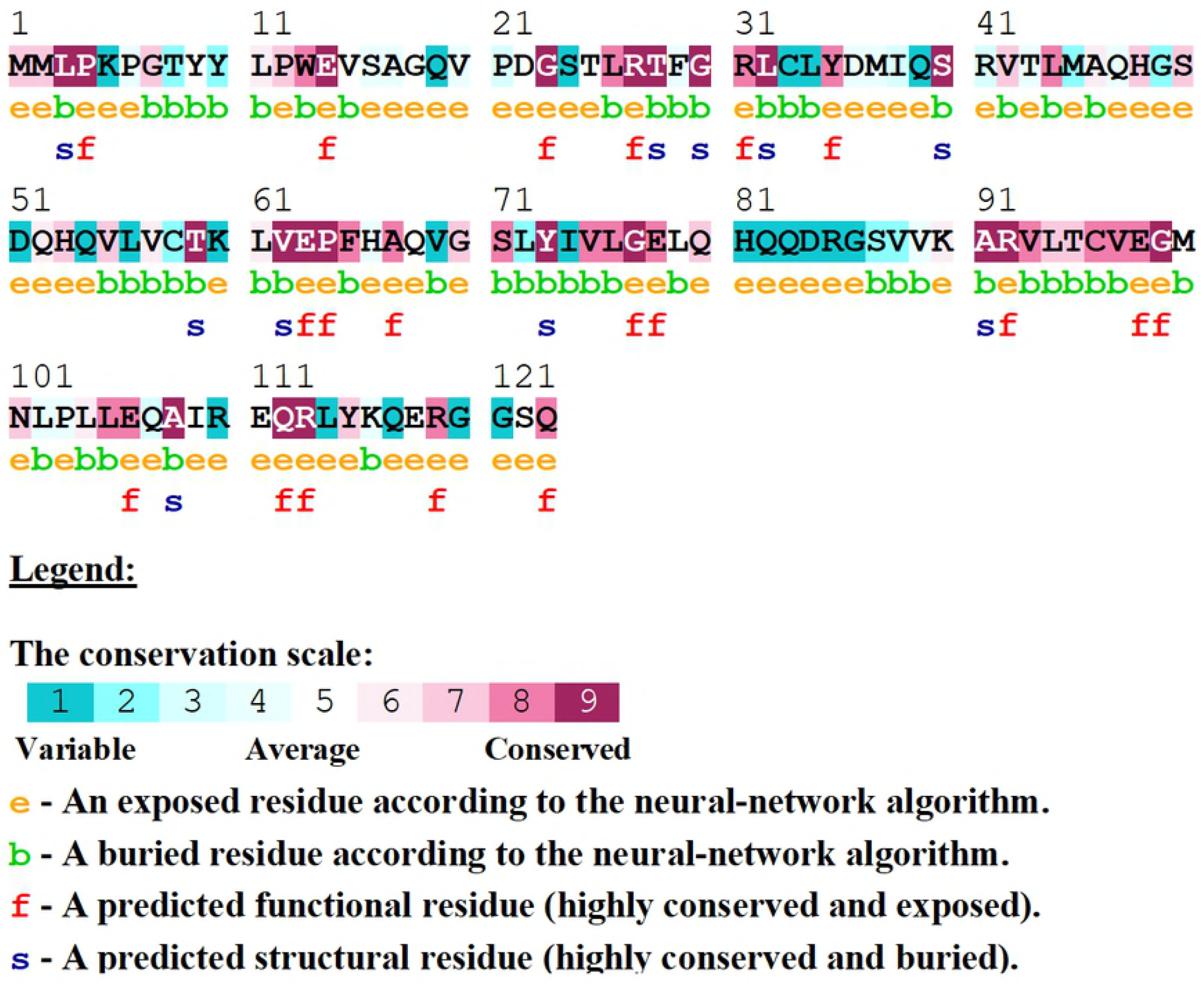
Conservation analysis of the TEN1 protein using ConSurf. ConSurf analysis also entails structural importance of a particular residue along with conservation score.

### Distribution of deleterious or destabilizing nsSNPs

TEN1 comprises of 123 amino acid residues and have one OB (oligonucleotide or oligosaccharides)-fold domain. The OB-folds domain was originally identified from a group of Yeast and bacteria [53]. OB-fold domain can bind and establish protein-DNA, protein-RNA, and protein-protein interactions [54,55]. Among these functions, interaction of OB-folds with ssDNA is extensively studied and characterized [11,56]. Structurally, the OB-folds are β-barrel consisting of five antiparallel β-strands capped by one α-helix at one end and has a binding cleft on the other end. The variability in length among OB-folds domain is mainly due to the differences in the lengths of variable loops connecting the conserved secondary structure elements [56].

Identification of relative percentage of high confidence nsSNPs in the OB-fold of TEN1 protein provides information about the relationship of a particular secondary structure component to be neutral or pathogenic. The secondary structure components; α1, β1, β2, β3, L3-4 (loop connecting the β3 and β4), β4, β5, α2, respectively have 75%, 60%, 28%, 50%, 45%, 22%, 77% and 35% deleterious or destabilizing mutations (**Fig. 5**). Mutations in the α1 and β5 are having more than 75% chance to be deleterious, while β1, β3 and L3-4 have about 50% chance. In addition, mutations in L1-2, L2-3, L4-5 and L5-α2 (loop connecting β5 and α2) suggested that nsSNPs occurring in these region has negligible chance to be deleterious. From these results, we can suggest that mutations in the α1, β1 and β5 are possibly more lethal than in other parts of TEN1. These observations were further complemented by sequence conservation analysis, which suggested that residues belonging to α1, β1 and β5 of TEN1 are highly conserved.

**Fig. 5:**
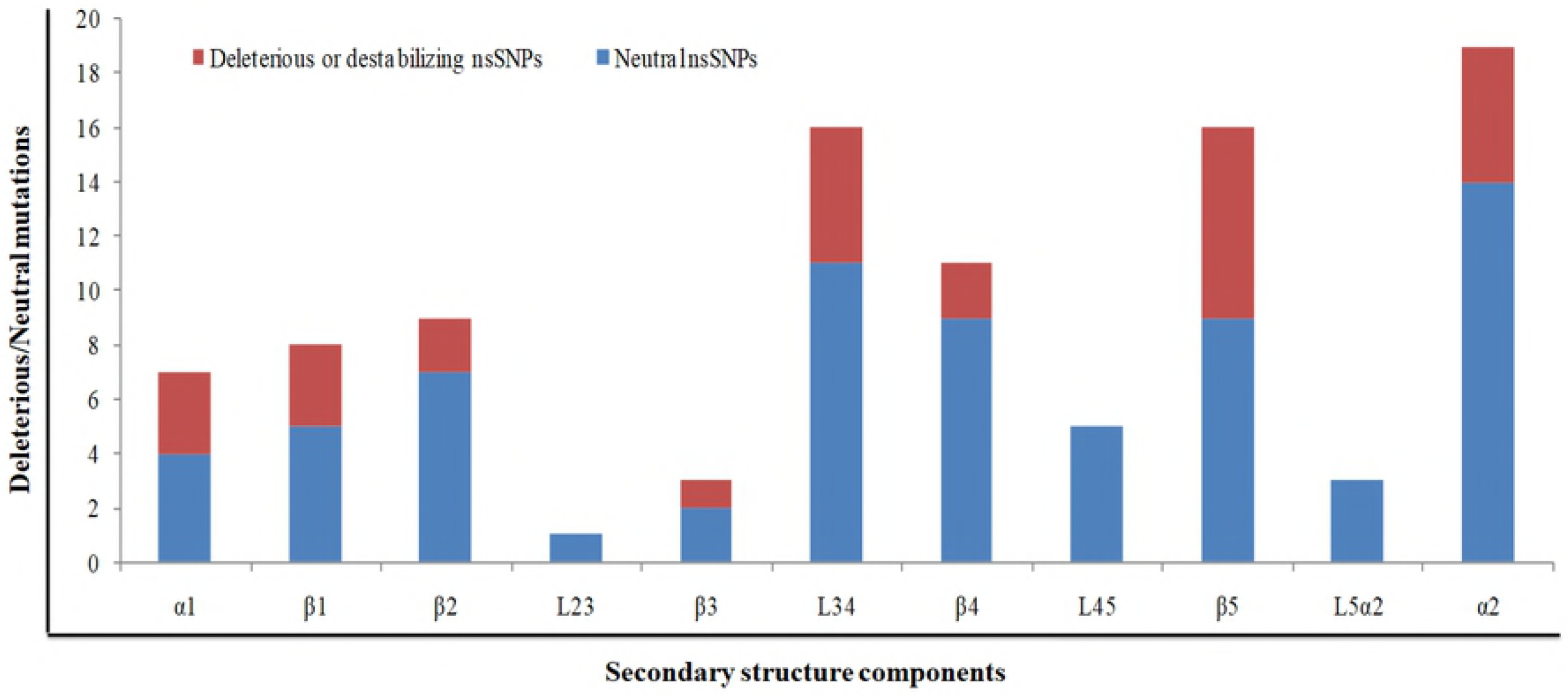
Distribution of deleterious/destabilizing and neutral nsSNPs in different structural components in TEN1 protein.

### Evaluation of disease phenotype

High confidence nsSNPs (deleterious and destabilizing) were analyzed for their phenotypic association using MutPred and PhD-SNP methods (**Table 1**). These methods predict a particular mutation as benign or pathogenic based on prediction score. MutPred and PhD-SNP methods depicts 58% (14) and 29% (10), respectively mutations are associated with disease phenotype. Of the 34 high confidence nsSNPs, we have identified only 8 missense mutations (W13G, L26P, C58Y, G70A, G77R, R92H, R92C and C96Y) which are reported as pathogenic from both methods. Pathogenic nsSNPs Trp13Gly found at the N-terminal flanking region, whereas Arg92His, Arg92Cys and Cys96Tyr are in the β5-strand.

**Table 1:** Prediction of disease phenotype analysis of high confidence nsSNPs in *TEN1* gene using PhD-SNP and MutPred prediction tools.

| S. No. | Variant ID | Variants | PhD-SNP | MutPred2 |  |
| --- | --- | --- | --- | --- | --- |
|  |  |  | Remark | Score | Remark |
| 1. | rs1322628164 | M2V | Neutral | 0.329 | Benign |
| 2. | rs892524367 | P4L | Neutral | 0.543 | Pathogenic |
| 3. | rs1212831970 | Y9C | Disease | 0.326 | Benign |
| 4. | rs1333358260 | W13G | Disease | 0.684 | Pathogenic |
| 5. | rs1224481693 | E14D | Neutral | 0.528 | Pathogenic |
| 6. | rs1175908725 | V15 F | Neutral | 0.584 | Pathogenic |
| 7. | rs1328038606 | G18V | Disease | 0.325 | Benign |
| 8. | rs964588646 | G23E | Neutral | 0.744 | Pathogenic |
| 9. | rs376979590 | T25M | Neutral | 0.171 | Benign |
| 10. | rs1262136645 | L26P | Disease | 0.855 | Pathogenic |
| 11. | rs1223059981 | D36N | Neutral | 0.301 | Benign |
| 12. | rs1250997925 | R41S | Neutral | 0.221 | Benign |
| 13. | rs1178755431 | L44V | Neutral | 0.286 | Benign |
| 14. | rs1412009927 | C58Y | Disease | 0.581 | Pathogenic |
| 15. | rs977512123 | L61M | Neutral | 0.168 | Benign |
| 16. | rs1032051988 | L61W | Neutral | 0.575 | Pathogenic |
| 17. | rs889310547 | P64T | Neutral | 0.510 | Pathogenic |
| 18. | rs951187486 | G70A | Disease | 0.482 | Pathogenic |
| 19. | rs1180274799 | G70S | Neutral | 0.545 | Pathogenic |
| 20. | rs1445270614 | Y73C | Neutral | 0.901 | Pathogenic |
| 21. | rs1358892195 | G77R | Disease | 0.880 | Pathogenic |
| 22. | rs562062613 | V88G | Neutral | 0.488 | Benign |
| 23. | rs1401886733 | A91V | Neutral | 0.831 | Pathogenic |
| 24. | rs1016457057 | R92H | Disease | 0.831 | Pathogenic |
| 25. | rs759839415 | R92C | Disease | 0.909 | Pathogenic |
| 26. | rs905216603 | V93M | Neutral | 0.543 | Pathogenic |
| 27. | rs1286634889 | C96Y | Disease | 0.922 | Pathogenic |
| 28. | rs1286634889 | C96F | Neutral | 0.906 | Pathogenic |
| 29. | rs368827427 | V97M | Neutral | 0.707 | Pathogenic |
| 30. | rs1216398771 | E106D | Neutral | 0.226 | Benign |
| 31. | rs1230794805 | R110W | Neutral | 0.130 | Benign |
| 32. | rs1158635929 | E111G | Neutral | 0.280 | Benign |
| 33. | rs772974788 | R119G | Neutral | 0.382 | Benign |
| 34. | rs772974788 | R119W | Neutral | 0.277 | Benign |

### Analysis of conformational changes in protein structure

Root mean square deviation (RMSD) is a commonly used quantitative measure of the similarity between two superimposed atomic coordinates, considered as a relative measure of structural and conformational changes in a given protein structure [57]. We have performed a comparative analysis of modeled tertiary structure of mutant proteins with the Wt to deduce possible structural and functional consequences imposed by pathogenic nsSNPs in TEN1 protein. We have superimposed the six pathogenic mutants (W13G, L26P, G77R, R92H, R92C and C96Y) of TEN1 protein onto structure of Wt protein using PyMol (**Figure 6A-F**). Mutation G77R in the β4-strand of TEN1 protein showed a remarkable conformational changes with a highest RMSD values in comparison to other mutations (**Figure 6C**). R92H and R92C mutations are involving the substitution of arginine by a small histidine and cysteine thus expected to affect the conformation of TEN1 protein which is evident from changes in RMSD values of backbone atoms (**Figure 6D-E**). Other three pathogenic mutations (W13G, L26P, and C96Y) are also showing a considerable structural change in the local structure as compared Wt.

**Fig. 6:**
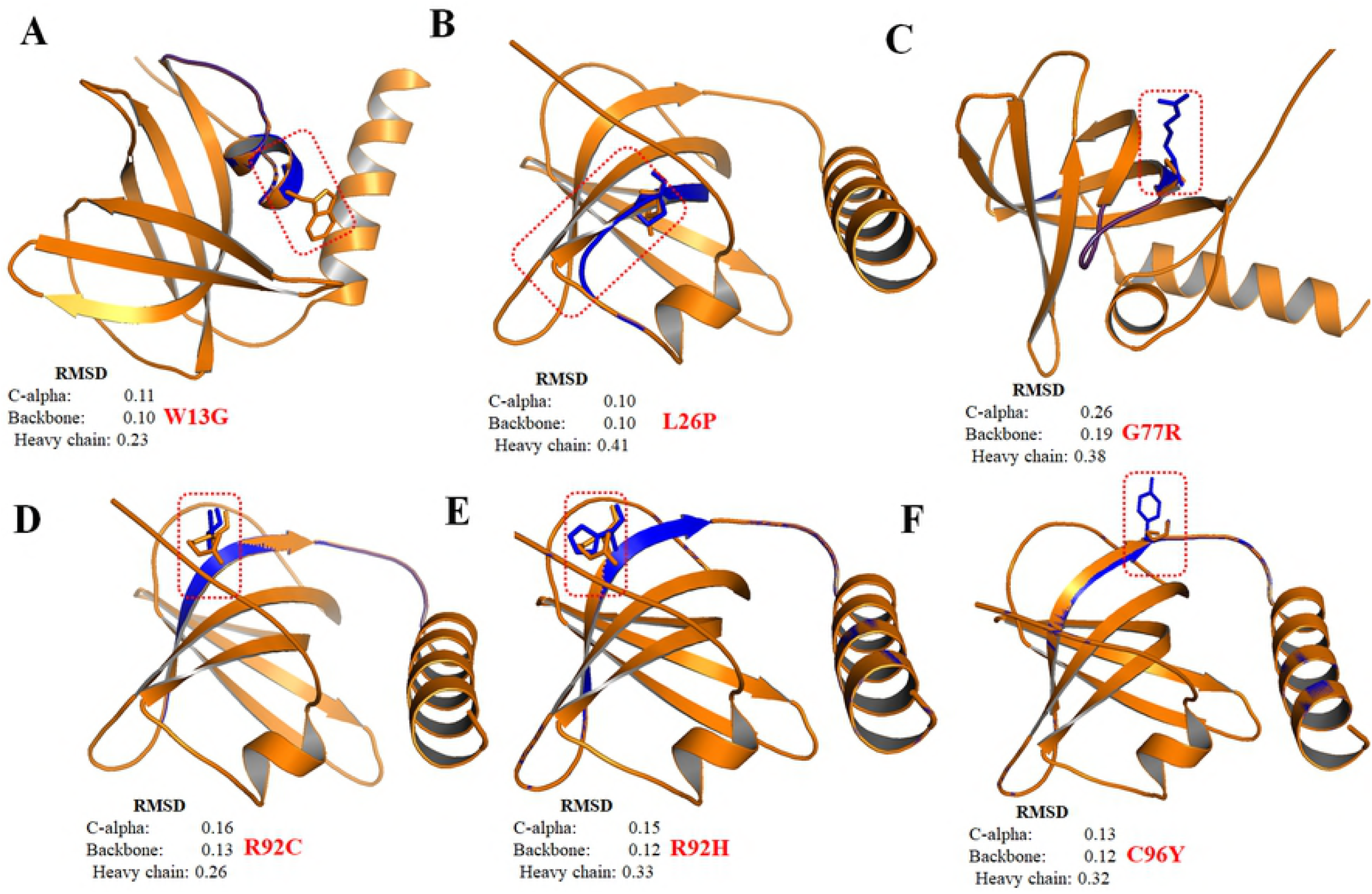
Structural superimposition of wild-type (Tan color) and mutant (Blue color) TEN1 proteins using PyMol. **(A)** W13G, **(B)** L26P, **(C)** G77R, **(D)** R92C, **(E)** R92H and **(F)** C96Y.

### Aggregation propensities analysis

Protein solubility is one of the critical attribute primarily related to its function [58,59]. Insoluble parts in proteins often tends to form aggregate which leads to development many disease including, amyloidoses [60], Alzheimer’s [61] and Parkinson diseases [62]. Aggregation propensity analysis was performed in the context of identification of disease or pathogenic SNPs. SODA classifies SNPs based on changes in α-helix and β-strand propensities; aggregation and disorder score, etc. Out of 8 pathogenic mutations obtained from MutPred and PhD-SNP tools, 6 (75%) were found to have increased tendency to form aggregate (**Table 2**). These aggregate forming potential of amino acid residue are primarily located at the C-terminal of TEN1 protein. Replacement of Arg92 by cysteine or histidine is considerably more prone to form aggregate in comparison to other pathogenic mutations.

**Table 2:** Predicted aggregation scores of wild-type and mutant TEN1 proteins using SODA server.

| Variants | Helix | Strand | Aggregation | Disorder | SODA | Remark |
| --- | --- | --- | --- | --- | --- | --- |
| 4JOI* | 0.293 | 0.316 | -4.44 | 0.089 |  |  |
| W13G | -0.211 | -0.75 | 4.7 | 0.748 | 4.072 | More soluble |
| L26P | -1.086 | -0.695 | 8.87 | 0.339 | 8.18 | More soluble |
| C58Y | -0.139 | 0.259 | -10.084 | -0.059 | -8.416 | Less soluble |
| G70A | 5.76 | -4.364 | -12.264 | 0.021 | -10.323 | Less soluble |
| G77R | 7.374 | -5.892 | -6.704 | 0.166 | -1.741 | Less soluble |
| R92H | 0.768 | -0.72 | -15.677 | -0.041 | -16.19 | Less soluble |
| R92C | 1.575 | -1.358 | -42.972 | 0.114 | -45.157 | Less soluble |
| C96Y | 1.861 | -1.429 | -8.643 | 0.067 | -6.102 | Less soluble |

4JOI\* = PDB ID of wild-type

### Structural and functional consequence of mutations

The OB-fold of TEN1comprises of five antiparallel β-strands folded into a complex β-barrel flanked by two α-helices (**Fig. 7**). N-terminal residues forming long coil and play crucial role in STN1-TEN1 complex formation. Following N-terminal coil, there is a short α-helix (α1) located at interface of two β-sheets known to provide stability to the structure. However, the C-terminal α-helix (α2) is situated at the opposite end of the β-barrel and spans whole length of the structure.

**Fig. 7:**
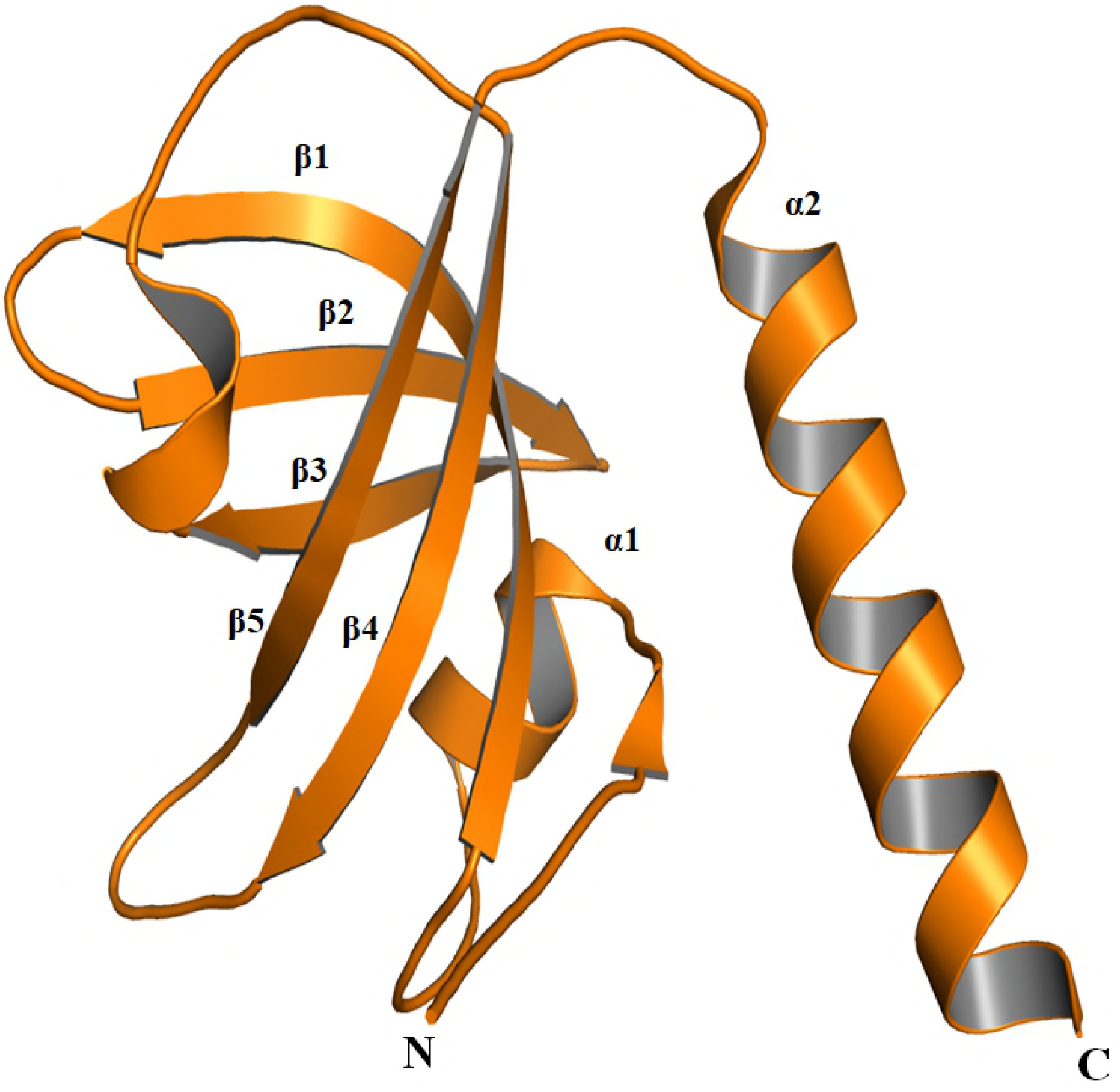
Cartoon representation of TEN1 protein (PDB ID: 4JOI).

The N-terminal of STN1 form stable heterodimer complex with TEN1. Complex formation between these two proteins are mediated by extensive interactions between the α2- and α3-helices of TEN1 and STN1, respectively (**Fig. S1A**). In addition to α-helices, β-barrels of TEN1 and STN1 also form extensive contacts (**Fig. S1B**) [25].

Some important amino acid residues including, Val159, Trp160, Ile164, Met167 and Leu168 of α3-helix and some region of flanking coils of STN1 form extensive hydrophobic contacts with the amino acid residues, Met100, Leu104, Leu105 and Ile109, of α2 of TEN1 (**Fig. S1C**). Additional interactions between the STN1 and TEN1 are mainly mediated by the conserved Tyr115 of TEN1 α2. Tyr115 is found at the interface of the STN1 and TEN1 and known to form extensive hydrophobic contacts with the side chains of Tyr49, Pro171 and Tyr174 of STN1. Similarly, interactions between the STN1 and TEN1 involve the surface of the β-barrels and the N-terminal tail of TEN1, that runs along the interface of the two domains and form extensive contacts with both of these two proteins (**Fig. S1D**). In particular, Arg27 of β1-strand and Arg119 of α2 of TEN1 make an important salt bridge with Asp78 of β2-strand and Asp33 of α2 of STN1, respectively. Further, Met167 of STN1 spans toward α2 and β-barrel interface of TEN1, and form extensive interactions with Leu105, Ala108 and Ile109 of α2 and Tyr9 of the N-terminal coil. It is fascinating that the STN1-TEN1 complex positions the ligand-binding pockets of each subunit on the same side of the heterodimer, forming an extensive ligand-binding pocket [25].

Mutations in protein is often coupled with destabilization or some time associated with disease pathogenesis. Previous studies on mutational analysis demonstrated that the effects of mutations on the stability of protein is primarily owing to changes in hydrophobic contacts [63,64,65].

However, subsequent studies in number of cases revealed that substitutions of large amino acid with smaller ones are usually accompanied by formation of cavity and effect residue depth and solvent accessibility [66,67,68,69]. To find out the impact of a particular mutation on the local and global environment of TEN1 protein structure, we have calculated van der Waals, hydrogen bonding, electrostatic and hydrophobic interactions in wt and mutant TEN1 using Arpeggio web server (**Table 3**) [70]. We have estimated change in the RSA, OSP and residue depth of wild-type and mutant TEN1 proteins (**Fig. S2**).

**Table 3:** Predictions of non-covalent interactions in WT and mutant TEN1 proteins using Arpeggio web server.

| **Variants** | **van der Waals Interactions** | **Hydrogen Bonds** | **Ionic interactions** | **Aromatic contacts** | **Hydrophobic contacts** |
| --- | --- | --- | --- | --- | --- |
| 4JOI* | 64 | 101 | 18 | 27 | 235 |
| P4L | 64 | 102 | 18 | 27 | 235 |
| W13G | 67 | 102 | 18 | 7 | 218 |
| E14D | 64 | 101 | 13 | 27 | 236 |
| V15F | 63 | 102 | 18 | 27 | 263 |
| G23E | 63 | 102 | 18 | 27 | 235 |
| L26P | 63 | 101 | 18 | 27 | 231 |
| C58Y | 64 | 101 | 18 | 27 | 242 |
| L61W | 64 | 103 | 18 | 27 | 240 |
| P64T | 64 | 102 | 18 | 27 | 235 |
| G70A | 64 | 102 | 18 | 27 | 235 |
| G70S | 64 | 102 | 18 | 27 | 235 |
| Y73C | 65 | 101 | 18 | 27 | 222 |
| G77R | 65 | 102 | 18 | 27 | 239 |
| A91V | 65 | 102 | 18 | 27 | 246 |
| R92H | 65 | 100 | 14 | 27 | 235 |
| R92C | 64 | 100 | 12 | 27 | 235 |
| V93M | 64 | 102 | 18 | 27 | 234 |
| C96Y | 65 | 101 | 18 | 35 | 237 |
| C96F | 65 | 101 | 18 | 28 | 243 |
| V97M | 64 | 102 | 18 | 27 | 240 |

4JOI\* = PDB ID of wild-type TEN1 protein.

Trp13 is a highly conserved and a buried residue of the N-terminal flanking coil and play important role in STN1-TEN1 complex formation. Substitutions of larger bulky and highly hydrophobic Trp13 by small, less hydrophobic glycine causes a subtle increase in the van der Waals and hydrogen bond interactions while a large decrease in stacking and hydrophobic interactions (**Table 1**). Differences in the size and polarity of substituted amino acid affecting the RSA, OSP and residue depth. Increased RSA value in the Trp13Gly substitution suggested that the substituted residue at Trp13 become more accessible to solvent, which is further supported by decrease in packing density (**Fig. S2**). Surface potential analysis shows a decrease in hydrophobicity in Trp13Gly substitution (**Fig. 8A**). The results suggested that substitution of Trp13 with the glycine seems indispensable for the stability of TEN1 structure.

**Fig. 8:**
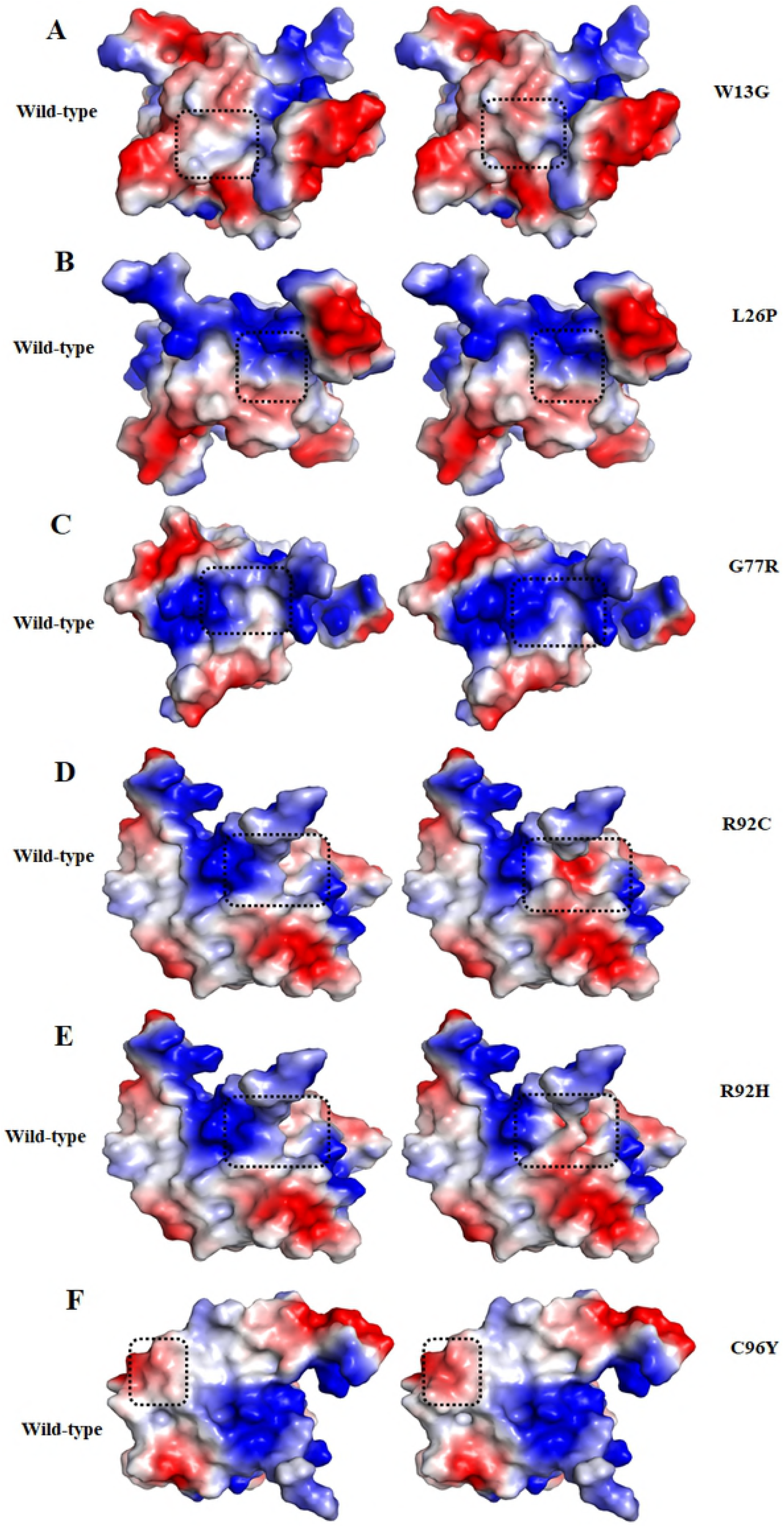
Surface potential representations of Wt (left panel) and mutant (right panel) TEN1 proteins. (A) **(A)** W13G, **(B)** L26P, **(C)** G77R, **(D)** R92C, **(E)** R92H and **(F)** C96Y. The color ramp for the electrostatic surface potential ranges from blue (most positive) to red (most negative). Surface potential of Wt and mutant residues are highlighted by dashed square.

Similarly, Leu26 is a highly conserved and buried residue found at the β1-strand of TEN1. Substitutions of hydrophobic Leu26 by a less hydrophobic proline effects only van der Waals and hydrophobic interaction at a little extent (**Table 1**). However, no any significant change was observed for RSA, OSP surface potential and residue depth by Leu26Pro mutation (**Fig. 8B**). We may conclude that the incorporation of imino group of proline may interfere with the folding pathway of TEN1 without effecting non-covalent interactions.

Gly77 is located in the β4-strand of TEN1, and plays important role in maintaining the structure and stability (**Fig. 7D**). Substitution of small, hydrophobic, highly conserved, exposed and functional Gly77 by a large and least hydrophobic, positively charged arginine shows an increase in the van der Waals, hydrogen bonding and hydrophobic interactions. In consistence, Gly77Arg mutation shows an increase in RSA, a subtle decrease in OSP and residue depth. Gly77Arg mutation increases positively charge environment in the vicinity of Gly77 (**Fig. 8C**). Lethality of Gly77Arg mutation is associated with the changes in RSA of surrounding residues which are critical to maintain the TEN1 stability.

Arg92 is belonging to the β4-strand of TEN1, and is important for the stability. Substitution of large, hydrophilic, highly conserved, exposed and functional Arg92 by a small hydrophobic, positively charged (histidine) and uncharged (cysteine) shows a disruption of one hydrogen bond and a large decrease in the ionic interactions. While, no significant change was observed in the van der Waals, stacking and hydrophobic interactions. Similarly, Arg92His and Arg92Cys mutations show an increase in RSA and a slight decrease in the OSP and residue depth. The increase in RSA suggesting that the substitution of Arg92 may increase the solvent accessibility of newly incorporated residues. A marked change in surface potential has been observed in Arg92His and Arg92Cys mutations (**Fig. 8D-E**). These results indicate the lethal effect of Arg92His and Arg92Cys mutations is primarily associated with the changes in hydrogen bonding, ionic interactions and RSA and thus protein stability.

Cys96 is situated in the β5-strand of TEN1. Substitution of small, less hydrophobic, highly conserved and buried Cys96 by a large and more hydrophobic tyrosine show an increase in the stacking and hydrophobic interactions. While no change was observed in other interactions. Cys96Tyr mutation shows an increase in the RSA and decrease in OSP. No significant change in surface potential has been found except an increase in the hydrophobicity (**Fig. 8F**). Our findings suggest that Cys96Tyr mutation may increases the important hydrophobic and stacking interactions which are being considered as a driving force for the protein stability. These increases in stability possibly overcome due to disruptions some important interaction Cys96.

## Conclusions

SNPs are considered as one of most recurring genetic variants associated with a number of diseases. Evaluation and understanding the role of mutations in different diseases are expected to shed light on disease susceptibility, and aid in the development towards more effective treatments. In this study, we have examined the consequences of nsSNPs in *TEN1* gene using advanced integrated bioinformatics approach. We have identified a large number of deleterious and destabilizing nsSNPs which are scattered in different secondary structural components of TEN1 with high chance of occurring in α1-helix and β5-strand. Aggregation propensities analysis of pathogenic mutation shows that 75% of pathogenic mutations in TEN1 have tendency to form aggregate and located at C-terminal of TEN1. In-depth structural analysis of these mutations reveals that the pathogenecity of these mutations may be driven through a large structural changes caused by loss/gain of non-covalent intramolecular interactions. The present study provides a mechanistic insights into the understanding pathogenic mutations in *TEN1* gene and their possible consequences.

